# Bacterial respiration during stationary phase induces intracellular damage that leads to dormancy

**DOI:** 10.1101/2020.11.27.401331

**Authors:** Spencer Cesar, Lisa Willis, Kerwyn Casey Huang

## Abstract

Most bacteria frequently encounter nutrient-depleted conditions, necessitating regulatory mechanisms that alter cellular physiology and allow for survival of starvation. Here, we show that regrowth of *Escherichia coli* from prolonged stationary phase upon encountering fresh nutrients is heterogeneous, with one subpopulation suddenly regrowing after a delay (dormancy) and another of nongrowing cells that represented an increasing fraction as the culture aged. Moreover, a sizeable fraction of cells rejuvenated immediately, even when the inoculum was from very old cultures. The size of the dormant and nongrowing subpopulations depended on the time cells had endured stationary phase, as opposed to time-dependent changes to the medium. Regrowth of dormant cells was correlated with the dissolution of polar phase-bright foci that likely represent aggregates of damage, and a deep-learning algorithm was able to distinguish cellular fates based on a single stationary-phase image. Growth restarted in dormant cells after the upregulation of chaperones and DNA repair enzymes, and deletion of the chaperone DnaK resulted in compromised stationary-phase cell morphology and higher incidence of non-growing cells. A mathematical model of damage accumulation and division-mediated partitioning was in quantitative agreement with experimental data, including the small population of cells capable of immediate regrowth even in old cultures. Cells that endured stationary-phase without the ability to respire all immediately and homogeneously regrew in fresh nutrients, indicating that respiration in stationary phase is the driver of dormancy. These findings establish the importance of intracellular damage control when nutrients are sparse, and repair when nutrients are plentiful.

## Introduction

Many bacteria in the environment live in nutrient-sparse conditions, motivating a deeper understanding of the physiological effects of starvation. As cells grow, they deplete the environment of nutrients and build up waste products that alter the environment. *Escherichia coli* cells sense and respond to these stressors through the sigma factor RpoS, which controls the expression of genes involved in acetate metabolism, pH homeostasis, and protection from oxidative stress during the initial stages of entry into stationary phase (1–3). Stationary-phase *E. coli* cells are typically thinner and shorter than log-phase cells due to successive reductive divisions upon entry into stationary phase (4), and cells can remain metabolically active long after cell expansion and division halts (5), suggesting that cellular physiology adapts during starvation.

The limited nutrient availability during stationary phase means that bacteria face particular challenges in maintaining homeostasis and viability. For example, genetic perturbations to the lipid transport machinery do not affect exponential growth but cause outer membrane vesiculation and cell lysis in stationary phase (6). Deletions of the super-oxide dismutases in *E. coli* result in massive cell death during stationary phase after growth in aerobic conditions (7), highlighting the importance of controlling reactive oxygen species. In rich media, after days in stationary phase, viability decreases by several orders of magnitude and a population of cells emerges with the “growth advantage in stationary phase” (GASP) phenotype, which results from successive sweeps of adaptive mutations with *rpoS* as a common target (8, 9). GASP cells can outcompete cells from earlier in stationary phase in starvation conditions, but have lower fitness during regrowth in fresh medium (8, 9), indicating tradeoffs between survival of stationary phase and rejuvenation.

The nutrient environment can greatly alter cellular survival in stationary phase and rejuvenation after starvation. During growth in minimal media, five days in stationary phase were required to reduce regrowth capacity and viability to levels comparable to LB after only one day in stationary phase (10). A theoretical model that postulated the slow accumulation of growth-inhibiting complexes during stationary phase predicted that mean lag time should scale with the amount of time spent in stationary phase and that the lag-time distribution should have a long tail (11). Indeed, a recent study showed that within a single stationary-phase population of *E. coli* cells, the distribution of single-cell lag times upon dilution into fresh medium can be highly heterogeneous, with a power-law tail that extends from 1 hour to >15 hours (12). In *Pseudomonas aeruginosa*, loss of multiple proteases accelerated cell death during growth arrest (13), indicating that active cellular processes maintain viability during starvation.

In addition to delayed regrowth and death, some starved cells enter a viable but nonculturable state, defined by prolonged metabolic activity but absence of growth for long periods of time (in some cases years) in permissible conditions (14). During aerobic growth, the non-culturable subpopulation of starved *E. coli* cells has more oxidative damage and higher levels of many genes and gene products related to oxidative stress (15), including chaperones such as DnaK and GroEL that repair misfolded proteins in protein aggregates (16, 17). The DnaK and GroEL regulator RpoH (18) is upregulated as cells transition from stationary to exponential phase before the first cell division (19), and transcriptional and post-translational positive feedback between RpoH and DnaK/GroEL (20) ensures high levels of chaperone expression when damaged proteins accumulate. However, how oxidative stress and chaperone production dictate the regrowth of single cells after starvation has yet to be determined.

Protein aggregates have been associated with growth-impaired bacterial cells in diverse contexts. After many generations of steady-state growth, aggregates accumulate at the older poles of aging *E. coli* cells, and are asymmetrically distributed after division (21). Cells with aggregates had reduced reproductive ability relative to their sister cells without aggregates (21), and new daughters had slightly faster growth rates than old daughters (22). Persisters are a clinically relevant subpopulation of non-growing cells defined by their survival of intense antibiotic treatment without resistance (23, 24). The formation of *E. coli* persisters in stationary phase was associated with protein aggregation (25), and in some cases persisters have been controversially connected to toxin-antitoxin systems that affect growth potential (26). Inhibition of respiration in stationary phase dramatically reduced the formation rate of persisters, which was correlated with cellular redox activity (27). Thus, aggregates appear to be a general sign of growth impairment, although the links to lag time in stationary phase and to redox activity remain unclear.

Here, we show that stationary-phase *E. coli* cultures develop into three distinct subpopulations based on their response to fresh nutrients: those that grow immediately, those with extended lag before regrowth (dormant), and those that do not resume growth at all. The fraction of dormant and non-growing cells increased over time, while >5% of cells persisted in being able to grow immediately even after 36 h of incubation. We show that growth deficiency is due to intracellular damage rather than waste buildup in the medium, and is correlated with the formation of large, polarly localized aggregates that dissipate upon the initiation of regrowth. Dormant cells increased their expression of chaperones and repair enzymes throughout lag time, and deletion of the chaperone DnaK disrupted cell shape in stationary phase and reduced regrowth capacity. A minimal model of the accumulation of damage and its asymmetric partitioning via cell division in stationary phase was in quantitative agreement with our experimental measurements and thus further supports the notion of damage-induced growth impairment. A pulse of fresh medium during stationary phase was sufficient to delay dormancy, further indicating that fresh nutrients enable damage repair. Growth-deficient subpopulations did not develop during anaerobic stationary-phase incubation in the absence of nitrate, highlighting the impact of respiration on dormancy.

## Results

### Culture age affects the heterogeneity of emergence from starvation

To reproducibly quantify when and to what extent cells enter dormancy (defined here as a state of delayed or inhibited growth) during stationary phase, we established a reproducible protocol involving growth of an *E. coli* culture inoculated from log-phase cells at OD_600_=0.1 (Methods). Starting at 12 h after inoculation, we extracted 1 μL of *E. coli* MG1655 cells every 2 h until 36 h, spotted them onto a 1% agarose LB pad, and acquired multi-hour time-lapse movies. To determine regrowth dynamics, we segmented trajectories of hundreds of cells from each movie and computed the instantaneous growth rate over time (Fig. 1A, Methods).

**Figure 1:**
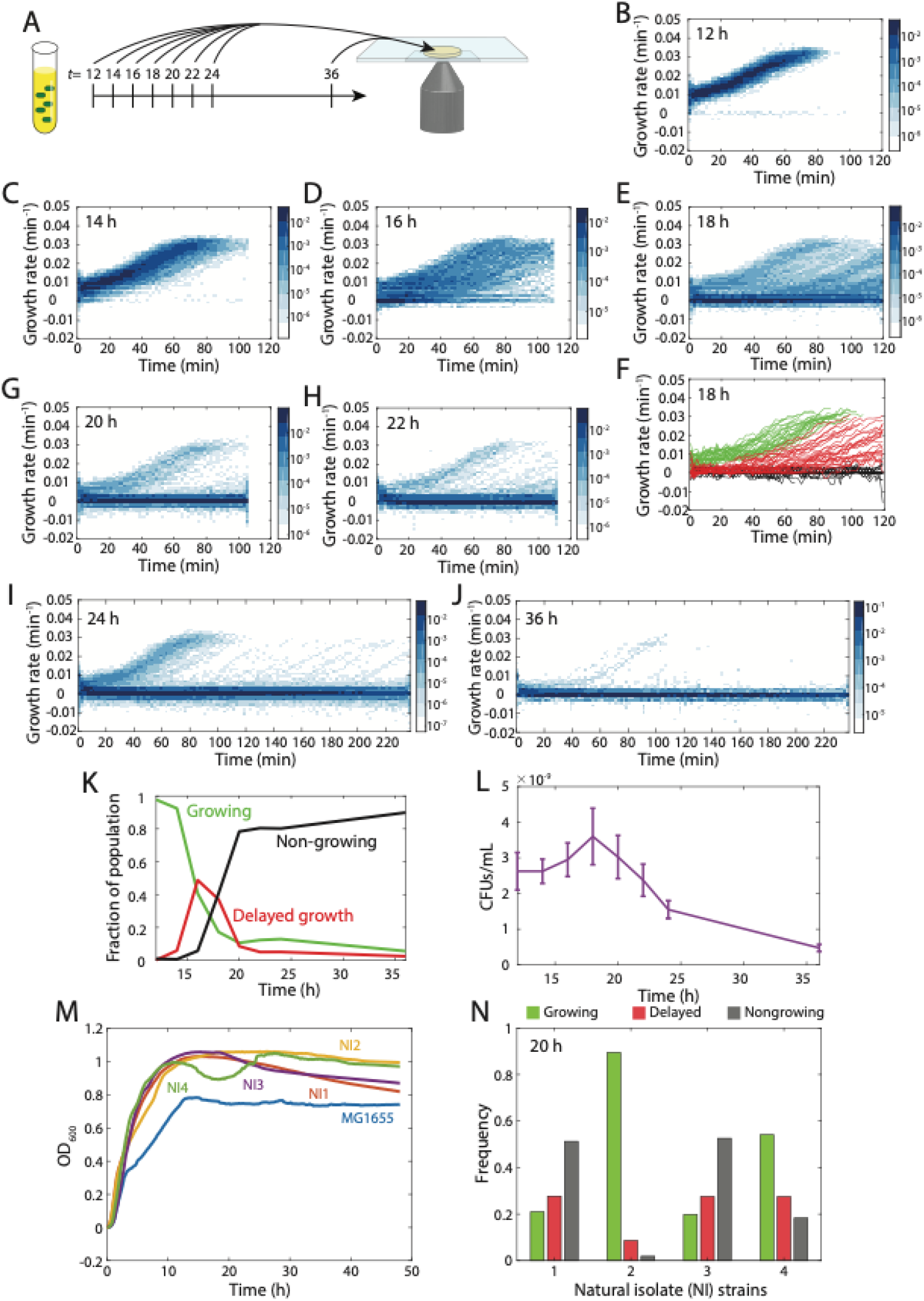
Increased time in stationary phase results in heterogeneous regrowth upon exposure to fresh medium. A) Schematic of protocol. Cells were sampled from a culture in stationary phase at the indicated times (in hours) and imaged on agarose pads with fresh LB to ascertain regrowth behaviors. B-E, G-J) Heatmaps of the distribution of instantaneous growth rates over time for cells placed on agarose pads made with fresh LB after 12 h (B), 14 h (C), 16 h (D), 18 h (E), 20 h (G), 22 h (H), 24 h (I), and 36 h (J). As the time in stationary phase increased to 16-18 h, the distribution of growth rates broadened and an increasing number of cells were non-growing. Past 18 h, the fraction of non-growing cells continued to increase. F) Single-cell growth trajectories after 18 h in stationary phase, classified as immediately growing (green), delayed growth (red), and non-growing (black) (Methods). K) The fraction of immediately growing cells began to decrease at 14 h but remained at 5-10% even after 36 h. The delayed-growth fraction peaked at 16 h and then decreased below the fraction of immediately growing cells. The non-growing fraction became the majority at ~18 h. L) The number of colony forming units (CFUs) increased slightly from 12 h to 18 h, suggesting a slow rate of division, and then steadily decreased thereafter. M) Four *E. coli* natural isolates all displayed higher yields than MG1655. N) After 20 h, the natural isolates exhibited distinct fractions of immediately growing, delayed-growth, and non-growing cells.

At 12 h, virtually all cells (>98%) resumed growth immediately and homogeneously, accelerating to their maximum growth rate of ~0.03 min^−1^ (doubling time of ~20 min) after ~60 min (Fig. 1B). However, by 14 h, some cells exhibited a delay in growth (Fig. 1C). By 16 h, the fraction of cells with delayed growth increased, and a small population of cells that did not grow for the entire period of imaging (henceforth referred to as “non-growing”) emerged (Fig. 1D). Cells with delayed growth typically had similar acceleration trajectories as growing cells once they began growth (Fig. S1A). At 18 h, the majority of cells did not resume growth while imaging (Fig. 1E). We identified thresholds based on the initial and final growth rates of each cell that robustly separated the three groups (immediate growth, delayed growth, and non-growing) for all culture ages (Methods, Fig. 1F).As the age of the culture increased, cells with delayed growth exhibited longer delays and increased heterogeneity (Fig. S1B-G) and the fraction of non-growing cells increased (Fig. 1E,G-J). Additionally, among the growing cells the initial growth rate upon exposure to fresh medium decreased with culture age (Fig. S1H). Nonetheless, cultures older than 24 h still maintained a small population of cells that grew immediately (Fig. 1I,J), suggesting they had not suffered ill effects of starvation.

Quantification of the prevalence of each subpopulation across culturing time revealed a transition from predominantly immediate growth at 12 h to a mixture of the three groups in which the delayed-growth fraction peaked at 16 h and then decreased with culture age (Fig. 1K). Notably, while the nongrowing fraction dominated after 20 h, the immediate-growth fraction was larger than the delayed-growth fraction beyond this point and continued to account for >5% even up to 36 h (Fig. 1K).

To determine whether the non-growing cells were dead, we plated cultures at each time point and counted colonies. CFUs/mL increased slightly from 12 to 18 h, suggesting the occurrence of cell division in stationary phase (Fig. 1L). After 18 h, CFUs/mL decreased steadily with culture age, with a ~2-fold and ~5-fold decrease relative to 12 h after 24 and 36 h (Fig. 1L). Overall, extended incubation in stationary phase induced heterogeneity in cellular recovery, including increased lag and eventually a transition to nongrowth and cell death.

### Natural isolates exhibit variable frequencies of dormancy and lysis

Laboratory strains such as MG1655 have been passaged in aerobic conditions for many generations, leading us to wonder if natural isolates of *E. coli* that generally reside in anaerobic environments within hosts would exhibit dormancy. We selected four strains (NI1-4) from a collection of *E. coli* clinical samples (28) and grew them along with MG1655 for 20 h, the incubation time for which MG1655 first exhibited a high level of dormancy (Fig. 1K). Somewhat surprisingly, all four natural isolates grew to a higher maximum OD than MG1655 (Fig. 1M). All four natural isolates exhibited dormancy, but with distinct fractions of each population. Interestingly, for the same time spent in stationary phase, all four natural isolates had fewer nongrowing cells than MG1655 (Fig. 1K). NI2 had the highest fraction of immediately growing cells (>80%), while NI3 had the lowest (<20%) (Fig. 1N). These data indicate that dormancy is generally conserved in *E. coli*, although with distinct kinetics across strains.

### Increased lag time is due to time spent in stationary phase and not changes in the extracellular environment

We considered two potential mechanisms for the cause of dormancy: intracellular accumulation of damage or toxins, or modification of the extracellular environment based on production of secreted toxic by-products or metabolites that affect cells directly or by modifying the environment. To distinguish between these two possibilities, we determined whether regrowth is driven by the time spent in stationary phase or by the age of the stationary-phase supernatant. We inoculated cultures 2 h apart, so that the ages of the two cultures were different. Once the culture ages had reached 10 and 12 h, we spun down each culture to separate cells from their supernatant (Methods). We then resuspended each pellet in the supernatants of the 10-h (“young”) and 12-h (“old”) cultures, and continued to incubate the resuspended cultures (Fig. 2A). Samples were extracted for imaging every 2 h thereafter, and we quantified the fraction of cells that exhibited regrowth after 1.5 h.

**Figure 2:**
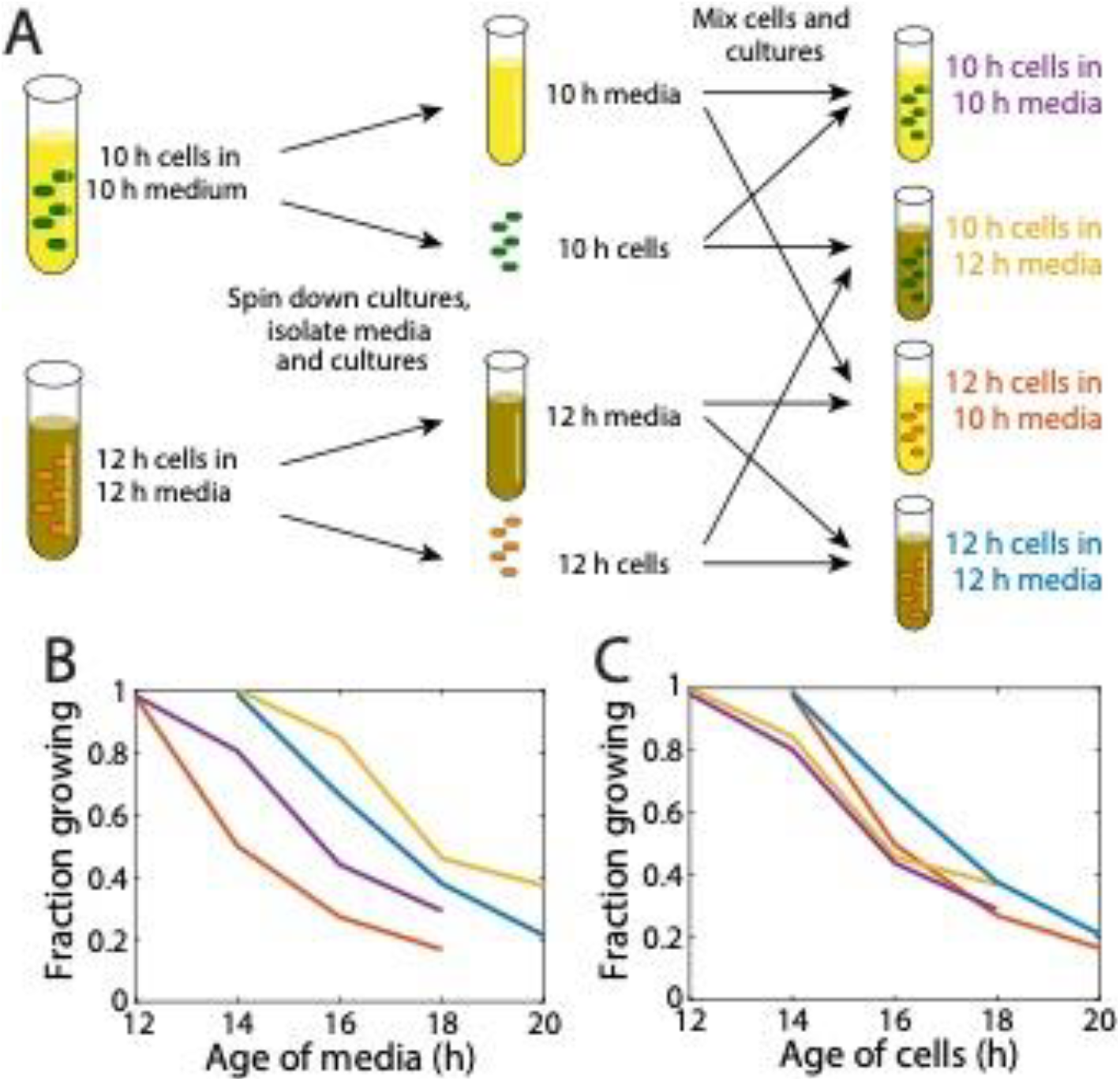
Dormancy results from changes to intracellular physiology rather than the extracellular environment. A) Scheme to distinguish the effects of time spent in stationary phase compared with modification of the extracellular environment. Overnight cultures diluted into fresh LB were grown for 10 and 12 h. Each was spun down and the supernatant was separated from the cells, whereupon all pairwise combinations of supernatant and cells were constructed and incubation of the four cultures continued. Samples were taken every 2 h thereafter and the fraction of growing cells on agarose pads with fresh LB was measured using time-lapse microscopy. B) The age of the culture supernatant does not dictate the onset of dormancy. Cells from a 12-h culture resuspended in 10-h supernatant transitioned to dormancy ~2 h later, while cells from an 10-h culture resuspended in 12-h supernatant transitioned to dormancy ~6 hours later. C) The dynamics of dormancy onset were highly similar for all four cultures in (B) when measured as a function of the time cells had been incubated.

Young cells in the young supernatant exhibited a transition from growing to nongrowing when the supernatant was ~16-h old, as did old cells in the old supernatant. Old cells resuspended in young supernatant transitioned to non-growth when the age of the supernatant was ~14 h (Fig. 2B), while young cells resuspended in old supernatant transitioned to nongrowth when the age of the supernatant was ~18 h (Fig. 2B). Notably, in all experiments the transition from growth to non-growth occurred ~16 h after inoculation. Indeed, the nongrowing fraction of all four combinations followed a common trajectory as a function of cell age (Fig. 2C). These findings suggest that dormancy is a result of accumulated physiological changes within cells, rather than changes to the supernatant.

### Visible protein aggregates form in late stationary phase and dissolve during regrowth

We noticed that cells from 18-h cultures, which had a high proportion of delayed and nongrowing cells (Fig. 1E), often had bright cytoplasmic foci visible in phase-contrast images (Fig. 3A,B). Previous studies of aging *E. coli* cells identified polar punctae of a YFP fusion to the small heat shock protein IbpA, which binds and colocalizes with protein aggregates (21). Another study observed aggregates associated with antibiotic persistence (23) that were composed of several metabolic, ribosomal, tRNA-related, and division-related proteins along with the stress-response sigma factor RpoS, and the aggregates dissipated when growth resumed (29). Thus, we hypothesized that the foci in stationary-phase cells represented aggregates that prevented immediate regrowth, such that cells with delayed growth would be visually distinct from those that could grow immediately. Indeed, phase-bright foci persisted in non-growing cells (Fig. 3A), dissolved from delayed-growth cells when growth restarted (Fig. 3B), and were absent from immediately growing cells (Fig. 3C). The foci were almost always located at poles and became more prevalent as the culture aged (Fig. 3D); the ramp up in the number of cells with 1 and 2 coincided with the increases in the nongrowing fraction (Fig. 1K) and the drop in viability (Fig. 1L), respectively. These data suggest that aggregates coincide with dormancy.

**Figure 3:**
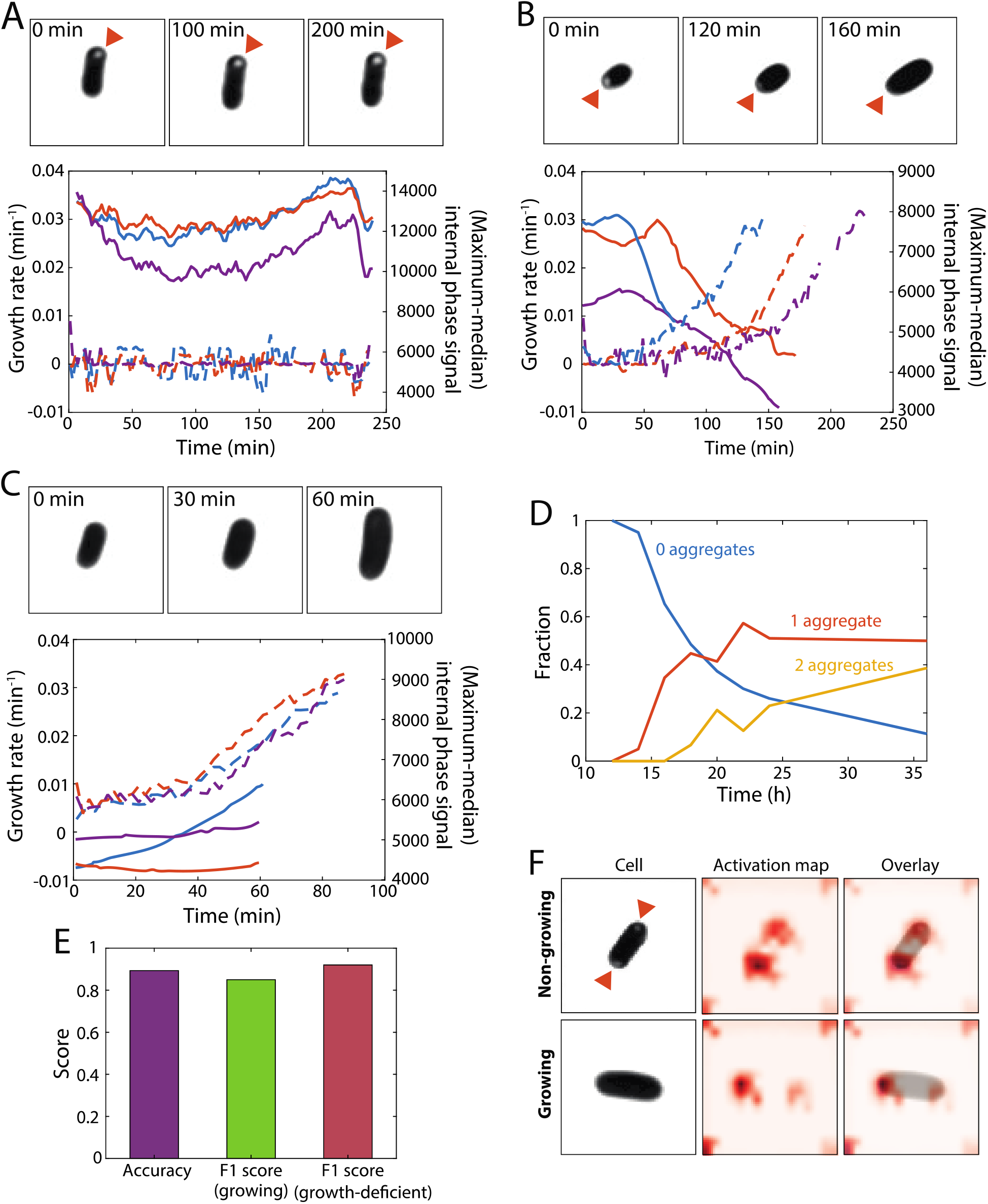
Delayed growth is associated with the dissolution of bright cytoplasmic foci that develop in stationary phase. A) Cells from a 20-h culture often exhibited visually apparent bright cytoplasmic foci (arrowheads) that were almost always located near one of the poles. Top: phase-bright foci persisted in non-growing cells. Bottom: the difference between the maximum and median phase signal inside the cell over time was used as an indicator of foci, and the intensities of foci in the cells shown remained approximately constant over time. Phase-contrast images were processed by contrast adjustment. B) Phase-bright dissolved from delayed-growth cells, coincident with the restart of growth, suggesting that the foci are connected with growth inhibition. Top: images were processed as in (A). Bottom: the intensities of foci in the cell shown started to decrease coincident with the growth rate increasing from zero. C) Phase-bright foci were absent from immediately growing cells. Top: images were processed as in (A). Bottom: the intensities of foci in the cells shown remained low over time. D) The number of cells with foci increased over time, and cells transitioned from predominantly having 0 foci to 1 or 2 foci similar to the transition to dormancy and non-growing (Fig. 1K). E) A deep-learning algorithm trained on phase-contrast images of cells from various cultures ages was able to accurately classify cells as immediately growing or growth-deficient (delayed/non-growing). The F1 score is the harmonic mean of the precision and recall. F) Activation maps demonstrate that the algorithm in (E) used information from the polar regions overlapping with foci to classify cells.

We trained a deep-learning algorithm on the first frame of phase-contrast time-lapse movies from cultures of various ages. For all movies, we used subsequent frames to classify cells as immediately growing or growth-deficient (delayed/non-growing) as before. Our algorithm was able to accurately classify ~90% of cells (none of which were used for algorithm training) with an F1 score of 0.8-0.9 (Fig. 3E), indicating that the future growth state of most cells is identifiable at the time of exposure to new nutrients. Activation maps showed that the classification algorithm focused on the polar regions of cells (Fig. 3F), consistent with the location of aggregates. These data suggest that aggregates are a general signature of growth inhibition during long-term starvation, and that dormancy is a state in which cells need restoration.

### Cells that take longer to emerge from stationary phase are smaller in size

Since aggregates were quantized and polarly localized, we speculated that the concentration of aggregates might be affected by cell size. We previously discovered that lag time during emergence from stationary phase for an *mreB*^*A53T*^ mutant with larger average cell width and volume was lower in glucose-supplemented minimal medium (30). To determine if regrowth was dependent on cell size in late stationary-phase cultures in LB, we computed the initial cell sizes of each cell (Methods) from the first frame of a time-lapse movie of a 20-h culture with a large fraction of nongrowing cells. We found that dormant cells were smaller than immediately growing cells, and non-growing cells were yet smaller (Fig. 4A).

**Figure 4:**
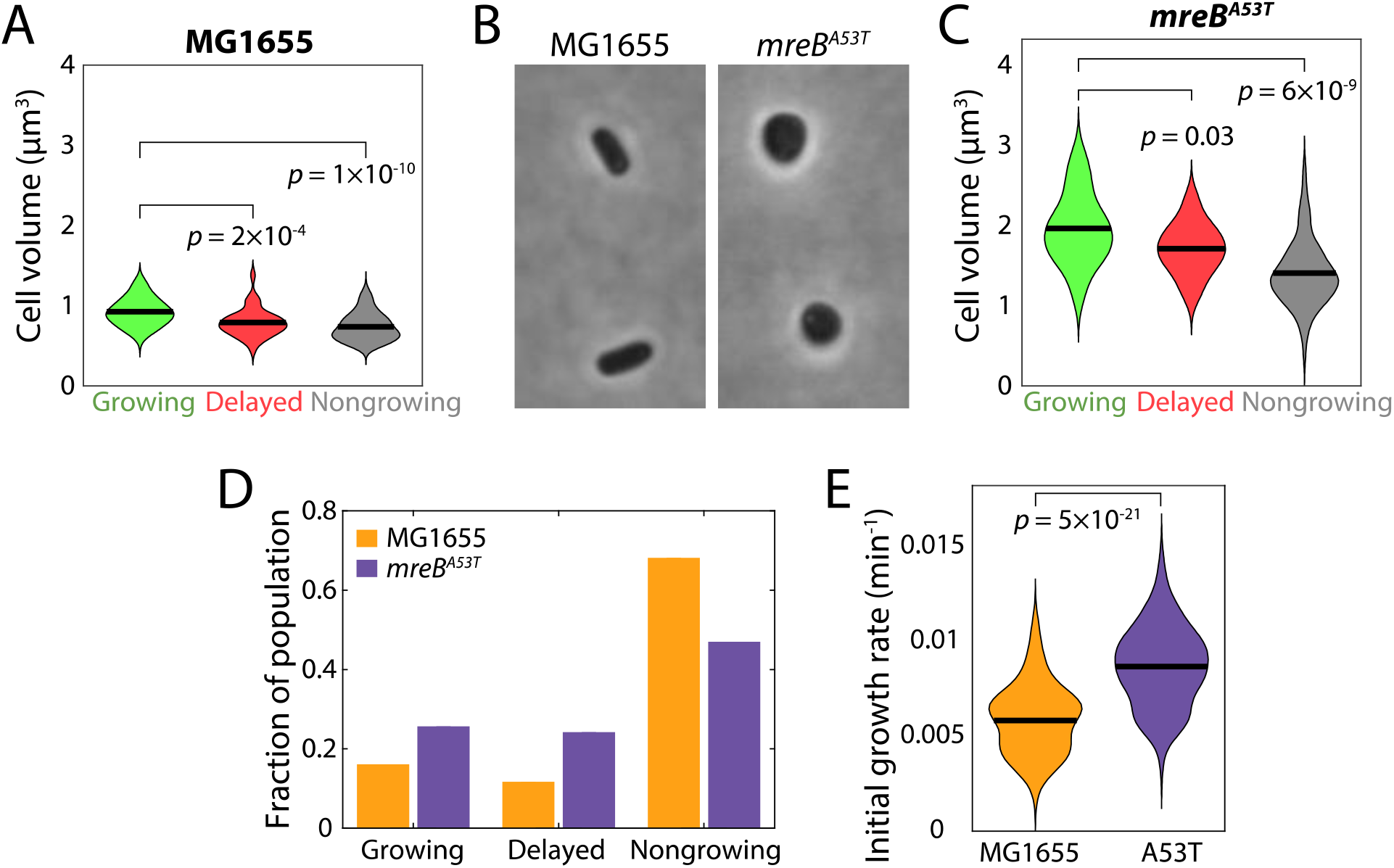
Cells with larger volume are enriched in the growing population and mutants with increased volume have higher growth potential during emergence from stationary phase. A) Delayed-growth cells from a 20-h culture were significantly smaller in the first frame of imaging than immediately growing cells, and non-growing cells were smaller yet. Thick horizontal lines are mean values. B) *mreB*^*A53T*^ cells from a 20-h culture were wider and larger in volume than wild-type MG1655 cells. C) *mreB*^*A53T*^ cells displayed a similar qualitative relationship between cell size and regrowth behavior as wild-type cells in (A), although *mreB*^*A53T*^ cells were larger overall. Thick horizontal lines are mean values. D) The *mreB*^*A53T*^ population in (C) had a larger fraction of immediately growing and delayed growth cells and a smaller fraction of nongrowing cells compared with wild type (A). E) The initial growth rates in the first 8 min of time-lapse imaging of immediately growing *mreB*^*A53T*^ cells were higher than those of wild-type cells, indicating that larger cells were generally more capable of stationary-phase exit. Thick horizontal lines are mean values.

To determine whether alteration of cell size via mutation was sufficient to alter regrowth capacity, we performed similar experiments with an MG1655 *mreB*^*A53T*^ mutant that has a larger volume than wild-type cells in stationary phase (Fig. 4A-C). In 20-h cultures, we found a similar qualitative relationship between cell size and regrowth behavior for the mutant as for wild-type (Fig. 4C). Moreover, there was a significantly larger fraction of immediately growing *mreB*^*A53T*^ cells and a smaller fraction of nongrowing cells than in a wild-type culture (Fig. 4D), and the initial growth rates of immediately growing MreB^A53T^ cells were faster than those of wild-type cells (Fig. 4E). Thus, cell size is associated with the ability of cells to emerge from stationary phase. However, the mean initial size of nongrowing *mreB*^*A53T*^ cells was larger than that of immediately growing wild-type cells, indicating that a threshold size does not dictate regrowth behavior.

### Delayed regrowth occurs after an increase in expression of chaperones and repair enzymes

A previous study suggested that non-culturable starved cells exhibit higher levels of several damage-associated stress regulons, including the transcripts *groEL* and *sodC* and the proteins RpoS and DnaK (15). To investigate whether these damage-associated regulons are also expressed in dormant cells, we performed stationary-phase regrowth experiments with *E. coli* MG1655 strains containing a GFP-reporter expressed from the promoter of candidate genes (here, these reporter strains are referred to as *P*<*gene*>-GFP) (31). We first focused on *dnaK*, which encodes a chaperone involved in DNA replication and repairing of protein aggregates (16, 17); DnaK was the strongest protein hit in the previous screen of non-culturable cells (15).

Time-lapse imaging of *PdnaK*-GFP cells from a 24-h culture on agarose with fresh LB revealed subpopulations of delayed-growth and non-growing cells, as we observed for wild-type. Initial GFP levels were similar for immediately growing, delayed-growth, and non-growing cells (Fig. 5A), and were uncorrelated with the time delay before rapid growth (Fig. 5B). Thus, stationary-phase *dnaK* expression is not predictive of dormancy. In immediately growing or non-growing *PdnaK*-GFP cells, GFP fluorescence remained approximately constant throughout imaging (Fig. 5C). Interestingly, in cells with delayed growth, GFP fluorescence increased substantially, indicating that expression of DnaK was specifically induced in these cells but only after the shift to fresh medium (Fig. 5C). Growth rate typically increased after an initial rise and plateau of *dnaK* expression (Fig. 5D), the fold-increase in *dnaK* expression was proportional to the delay before rapid growth (Fig. 5E), and in all but the cells with the shortest delays the maximum in fluorescence occurred at approximately the same time as growth resumption (Fig. 5F). These data suggest that cells with longer lag times had higher levels of damage and hence required more chaperone activity.

**Figure 5:**
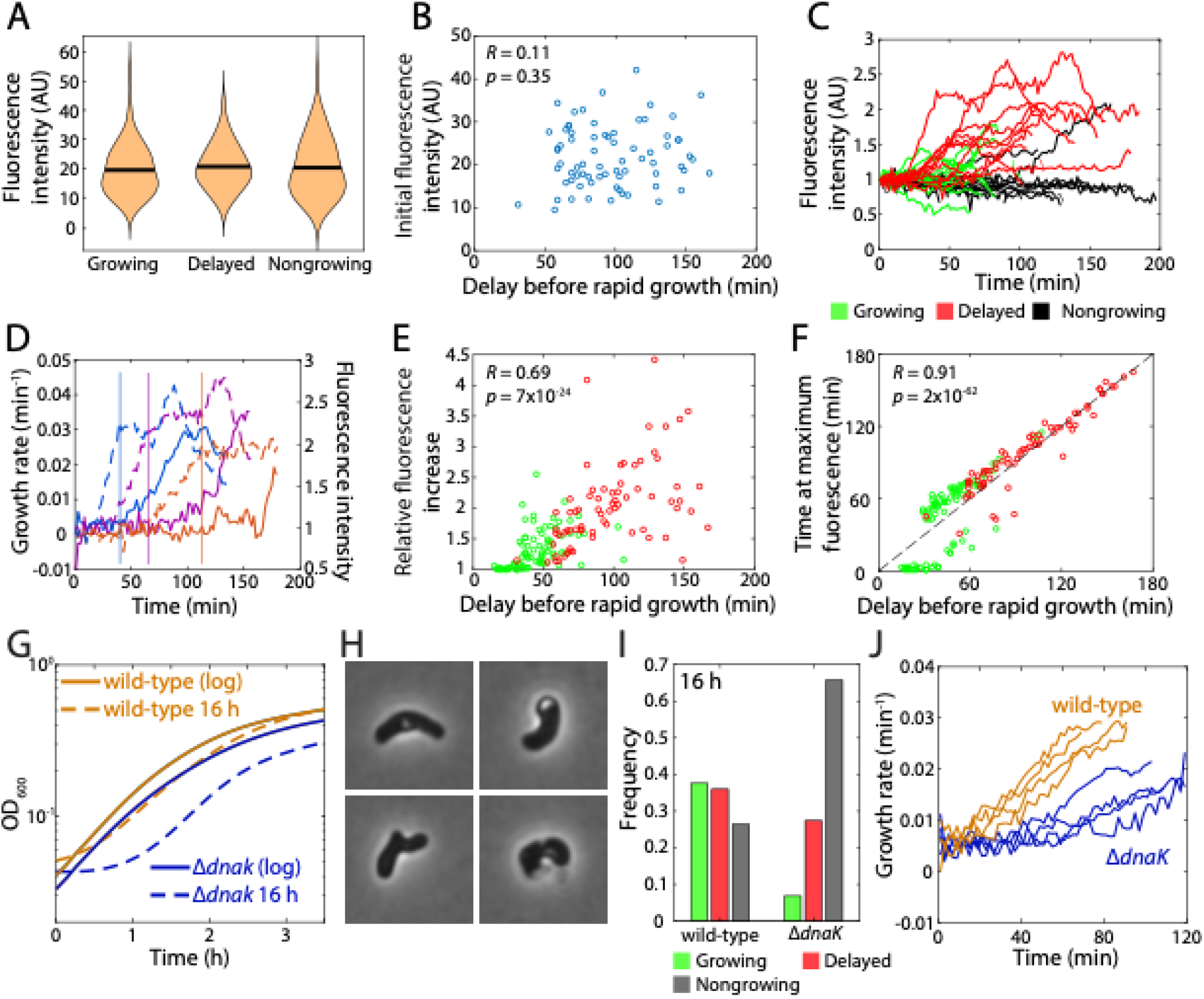
Chaperone activity is important for the revival of dormant cells and for survival of stationary phase. A) Initial fluorescence intensity in stationary-phase cells with a *gfp* reporter expressed from the *dnaK* promoter was similar for cells in all regrowth groupings. B) Among delayed-growth cells, the initial GFP intensity was uncorrelated with the delay before rapid growth (>0.02 min^−1^), indicating that *dnaK* expression in stationary phase is not a predictor of regrowth behavior. C) In delayed-growth cells (red), GFP intensity (normalized by initial fluorescence) increased, while in immediately growing (green) or non-growing (black) cells, intensity remained approximately constant over time. D) In delayed-growth cells, GFP intensity (dashed curves) (normalized by initial fluorescence) increased up to a plateau before the instantaneous growth rate (solid curves) started to increase from zero. E) In delayed-growth and immediately growing cells, the maximum increase in GFP intensity relative to the initial value from stationary phase was highly correlated with the delay before rapid growth. F) For most of the cells in (E), the time at which the maximum fluorescence was reached was approximately coincident with the delay before rapid growth was achieved. Dashed line is *y*=*x*. G) Growth curves of BW25113 wild-type and Δ*dnaK* cultures started from log phase were similar, while the Δ*dnaK* growth curve started from stationary phase exhibited increased lag. H) Δ*dnaK* cells from a 16-h culture exhibited aberrant morphologies compared with the short rods typical of a wild-type culture. I) A significantly higher fraction of Δ*dnaK* cells compared with wild-type cells were non-growing after 16 h, and very few grew immediately. J) Of the small population of immediately growing Δ*dnaK* cells, the initial growth rate was significantly lower than wild-type cells.

We observed behavior quantitatively similar to DnaK in *PrecA*-GFP (RecA is a DNA repair enzyme) (Fig. S2A) and *PgroE*-GFP (GroEL is a chaperone) cells (Fig. S2B), further supporting the notion that dormancy is due to accumulated DNA and protein damage. By contrast, for several non-damage-related promoters including *ompA* (which encodes an abundant outer membrane porin), *pyfK* (pyruvate kinase), *rpsB* (a ribosomal protein subunit), *tnaC* (tryptophan operon leader peptide), and *tufA* (translation elongation factor Tu 1), GFP increases were not correlated with the delay until rapid growth (Fig. S2C-G), arguing against the need to boot up metabolic, translational, or cell-envelope synthesis capacity.

To determine whether DnaK is critical for regrowth from stationary phase, we examined the Δ*dnaK* knockout from the Keio collection (32) and its parent strain BW25113. Growth curves of BW25113 wild-type and Δ*dnaK* cultures inoculated from log-phase cells at OD_600_=0.1 were similar (Fig. 5G), indicating that the mutant can grow comparably to wild-type in exponential phase. However, Δ*dnaK* cultures inoculated from stationary phase exhibited longer lag than wild-type (Fig. 5G), suggesting that growth defects emerge in stationary phase. Indeed, after 16 h of incubation, Δ*dnaK* cells exhibited distorted morphologies (Fig. 5H) in stark contrast with the typical short rods of wild-type cells. Whereas most wild-type BW25113 cells re-grew immediately or exhibited delayed growth, ~70% of Δ*dnaK* cells were nongrowing (Fig. 5I). Of the small subpopulation of immediately growing cells, their growth rate kinetics were substantially slower than the immediately growing population of wild-type cells (Fig. 5J). Moreover, 15-20% exhibited a range of growth-impaired behaviors not observed in wild-type cells, including growth for only a short period upon transition to fresh LB as well as cycles of expansion and shrinking (Movie S1). These data confirm the pivotal role of DnaK in surviving stationary phase and in resuming growth in fresh medium, presumably by repairing damage that accumulates during starvation.

### Mathematical model quantitatively predicts dormancy kinetics

Our imaging strongly suggests a link between regrowth of dormant cells and the dissolution of phase-bright foci (Fig. 3), but so far causality has not been established. Thus, we sought to test whether our findings are quantitatively consistent with a model of damage-mediated growth inhibition. The increased fraction of non-growing cells in older cultures (Fig. 1K) suggested that damage is ongoing throughout stationary phase. Since aggregates appeared primarily at the poles (Fig. 3A,B), we surmised that division of a cell with only one aggregate would segregate most damage to a single daughter cell, restoring the other daughter to a (nearly) undamaged state, similar to previous findings of the rejuvenation of older cells via asymmetric aggregate segregation after division (21). The increase in CFUs/mL from 12 to 18 h (Fig. 1L) suggested that at least some cells are capable of continued growth and/or division in stationary phase, albeit slowly due to nutrient limitation.

In our model, we assume that damage accumulates spontaneously throughout stationary phase, and high levels of damage cause a delay in regrowth and eventually lead to a non-growing state. Damage accumulates from 0 upon entry into stationary phase (which we estimate at ~11 h after inoculation) probabilistically at a rate δ. Cells with damage <*A*_1_ grow immediately upon restoration to fresh medium, cells with damage between *A*_1_ and *A*_2_ experience delayed growth, and cells with damage >*A*_2_ are assumed to be non-growing (at least during the period of observation; Fig. 6A). Simulations showed that cells transit from immediate to delayed growth on average at time 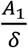 and then to non-growth at 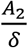, with a peak in the fraction of delayed-growth cells of magnitude 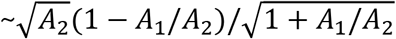 (Fig. 6B). For 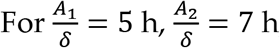, and *A*_2_ = 20, the model recapitulated regrowth statistics up to, but not beyond, ~20 h after inoculation (Fig. 6B, S3A).

**Figure 6:**
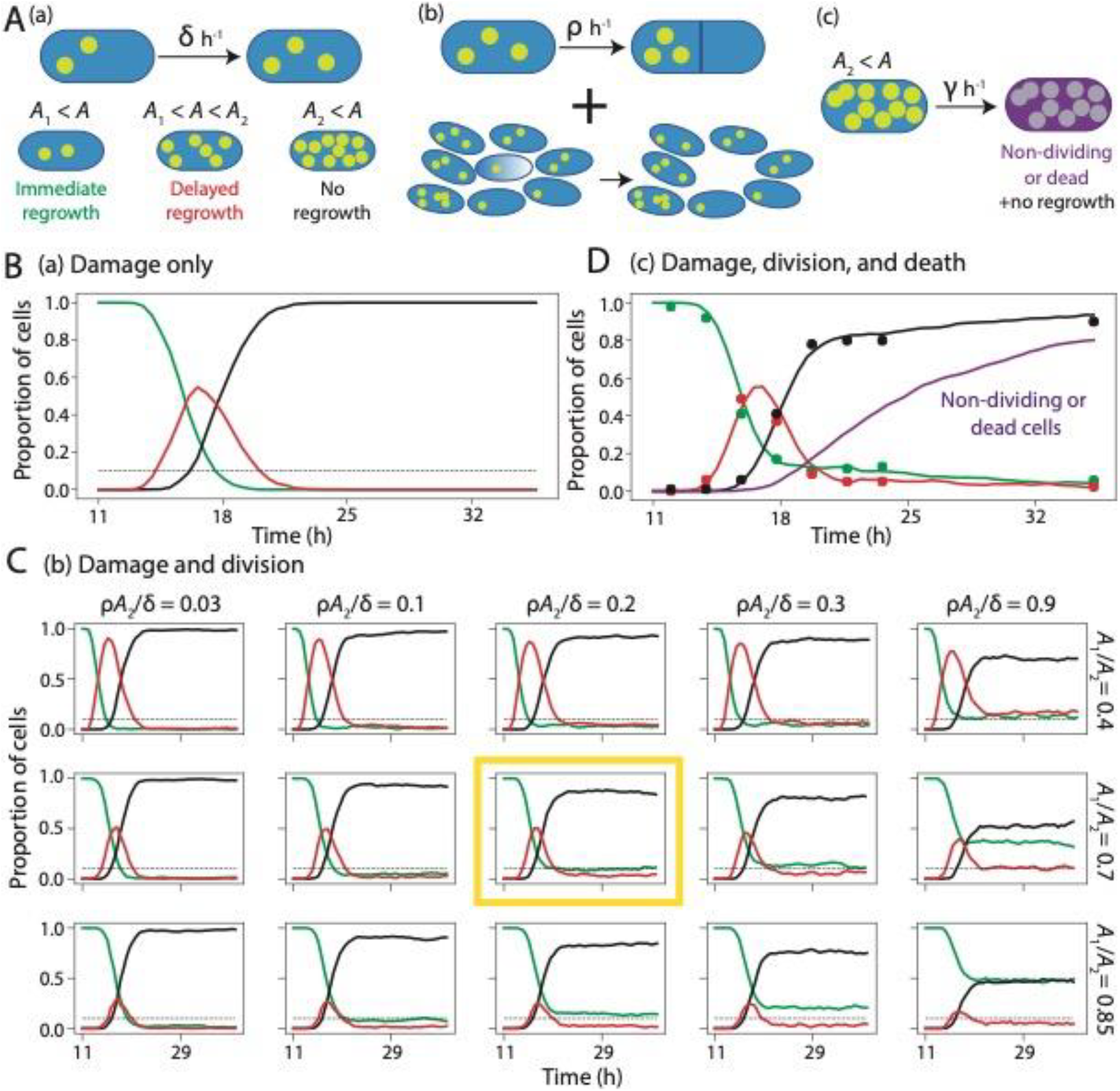
A model of stationary-phase damage accumulation recapitulates dormancy kinetics when lag time is damage-dependent and a subset of cells divide slowly. A) (a) In a model with only damage, throughout stationary phase starting ~11 h after inoculation, damage *A* accumulates at rate δ h^−1^. Upon restoration to fresh medium, cells grow immediately if *A* ≤ *A*_1_, grow with a delay if *A*_1_ < *A* ≤ *A*_2_, and do not grow during the period of observation if *A* > *A*_2_. (b) Division is incorporated at rate *ρ* h^−1^. Upon division, damage partitioning is asymmetric such that one daughter receives all the damage. (c) Cells with *A* > *A*_2_ stop dividing or die (without lysis) at rate *γ* h^−1^, such that they do not form colonies upon restoration to fresh medium during CFU counting. B) Simulations based on model (a) with damage only using 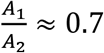 and *A*_2_ ≈ 20 recapitulated the early fractions of immediately growing (green), delayed (red), and non-growing (black) subpopulations. However, the persistent subpopulations with immediate and delayed regrowth after ~20 h did not occur. The horizontal dashed line represents the experimental estimate of the fraction of immediately regrowing cells at 24 h of 0.1. C) The addition of division with asymmetric-damage partitioning (model (b)) with 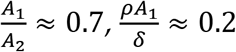, and *A*_2_ = 20 recapitulated the early fractions and the long-term persistent subpopulations of immediate and delayed regrowth (yellow box). Deviations from these parameter values altered the dynamics. D) Simulations in which cells become non-culturable as damage accumulates (model (c)) produced excellent fits to experimental data (circles, Fig. 1K) for 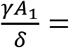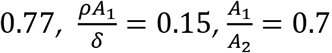, and *A*_2_ = 25. By 36 h after inoculation, ~80% of cells were predicted to be nonculturable, consistent with CFU measurements (Fig. 1L).

Without the potential for cellular recovery, the model predicts a negligible proportion of immediately growing cells after ~20 h (Fig. 6B), coincident with the transition to non-growth (because cells only become progressively more damaged), which is inconsistent with the observed persistence of an immediately growing population (Fig. 1K). Thus, we incorporated into our model division at a probabilistic rate *ρ*. Division restores one daughter cell into an undamaged (and thus immediately growing) state, while the other daughter inherits all damage (Fig. 6A). At long times, the proportion of immediately growing cells is 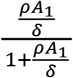 and the ratio of immediately growing to delayed growth cells is 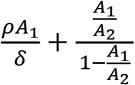 (SI, Fig. S3B). Our experimental measurements indicate that 36 h after inoculation (Fig. 1K), 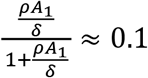 and 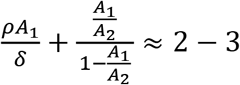, hence 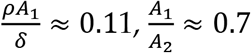, and the average number of divisions over the time it takes a cell to reach a non-growing state is 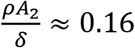. Remarkably, simulations with these parameter values (Fig. 6C) recapitulated both early and late stationary-phase regrowth statistics (Fig. 1K), while significant deviations from these values resulted in poor fits to the data (Fig. 6C), regardless of whether absolute levels or concentrations of intracellular damage trigger dormancy (Fig. S3C). The predicted time between divisions is 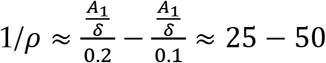h. Results were similar if the division rate declines linearly with increased damage (SI, Fig. S3D), which is plausible since FtsZ is sequestered in aggregates at the cell poles in stationary phase (Fig. S4) (29).

Viability declined starting ~18 h after inoculation (Fig. 1L), approximately coincident with the transition to non-growth. We assessed the ability of our model (Fig. 6A) to account for the number of cells in a non-viable state late in stationary phase by including a transition to non-viability at a rate *γ* after accumulating *A*_2_ of damage. For 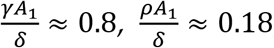, and 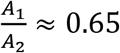, our model predicted that ~80% of cells become nonviable by 36 h (Fig. 6D), similar to our experimental data (Fig. 1L). In sum, a minimal model containing reasonable assumptions quantitatively explains the trajectories of immediate-growth, delayed-growth, non-growing, and non-viable cells as a function of culture age, suggesting that a single variable that accumulates over time (i.e., damage) and is asymmetrically partitioned by cell division is sufficient to explain dormancy kinetics.

### A pulse of fresh medium delays the onset of dormancy

To confirm that intracellular damage built up during starvation is reversible, we administered a pulse of fresh medium at a time point when many cells were entering a dormant state and quantified its effects on regrowth kinetics. We grew dilutions of two overnight cultures for 12 h, and then subjected the cultures to separate treatments. One was spun down and resuspended in 2X its volume of fresh medium. The second was spun down and resuspended in its own spent supernatant.

We incubated both cultures for 20 min, resuspended them in spent LB from a 12 h-old culture, and incubated them for a further 10 h. We extracted samples every 2 hours, and imaged cells on agarose pads with fresh LB to examine their emergence from stationary phase. For both cultures, most cells were able to grow immediately at 12 h (Fig. S5). The second culture (no pulse of fresh medium) behaved as before, with a gradual increase in the fraction of nongrowing cells such that ~50% were nongrowing 4-6 h later (Fig. S5). By contrast, the 20-min pulse of excess fresh medium was sufficient to ensure that after 4 h back in spent medium almost all cells retained the ability to resume growth immediately on fresh medium (Fig. S5). Nonetheless, a similar fraction of cells were non-growing (~80%) in all cultures after 10 h (Fig. 2D), suggesting that the pulse reversed or delayed damage but only temporarily. The fact that the pulse had an effect even though it was administered before any cells entered dormancy is consistent with the accumulation of damage prior to growth impairment.

### Cells do not enter dormancy in the absence of respiration

Inhibition of respiration in stationary phase reduces the formation of persisters (27), and protein aggregation during stationary phase is oxygen-dependent (33). Thus, we hypothesized that reactive oxygen species are the perpetrators of stationary phase-acquired damage. To test this hypothesis, we first grew a culture in non-shaking conditions, which allows cells to rapidly deplete oxygen (34). Now, after 20 h, most cells were able to grow immediately upon exposure to fresh medium (Fig. S6A,B), unlike in shaking conditions (Fig. 1K).

To determine whether growth without oxygen prevents dormancy, we grew MG1655 cells in an anaerobic chamber using pre-reduced LB (Methods). Periodically, a sample of cells was extracted for imaging in an aerobic environment. We observed a stark contrast with cells grown aerobically: in the anaerobic chamber, even 40 h after inoculation, all cells immediately resumed growth on fresh LB (Fig. S6C), suggesting that respiration was the cause of dormancy.

To test whether oxygen exposure specifically during stationary phase was responsible for dormancy, and to control for the different yields (Fig. S6D) and extracellular environments of *E. coli* cultures grown aerobically and anaerobically, we grew a culture aerobically for 12 h and then resuspended it in supernatant from an 18-h aerobic culture (which was fully saturated) that had been pre-reduced in the anaerobic chamber (Fig. 7A, Methods). We then incubated the resuspended culture in the anaerobic chamber for a further 36 h. Similar to our observations involving cultures grown entirely within the anaerobic chamber (Fig. S6C), all cells grew immediately (Fig. 7B). Moreover, no cells (of >100 examined) displayed any sign of aggregates, suggesting that damage was minimal (or nonexistent). Finally, to determine if it was the absence of oxygen or the incapacity to respire that protected cells during anaerobic growth in LB, we grew cells anaerobically in LB+10 mM nitrate to allow for anaerobic respiration. Now, delayed and nongrowing populations emerged after 48 h, with only ~40% of cells able to immediately resume growth (Fig. S6E). Thus, respiration during stationary phase is a primary cause of dormancy.

**Figure 7:**
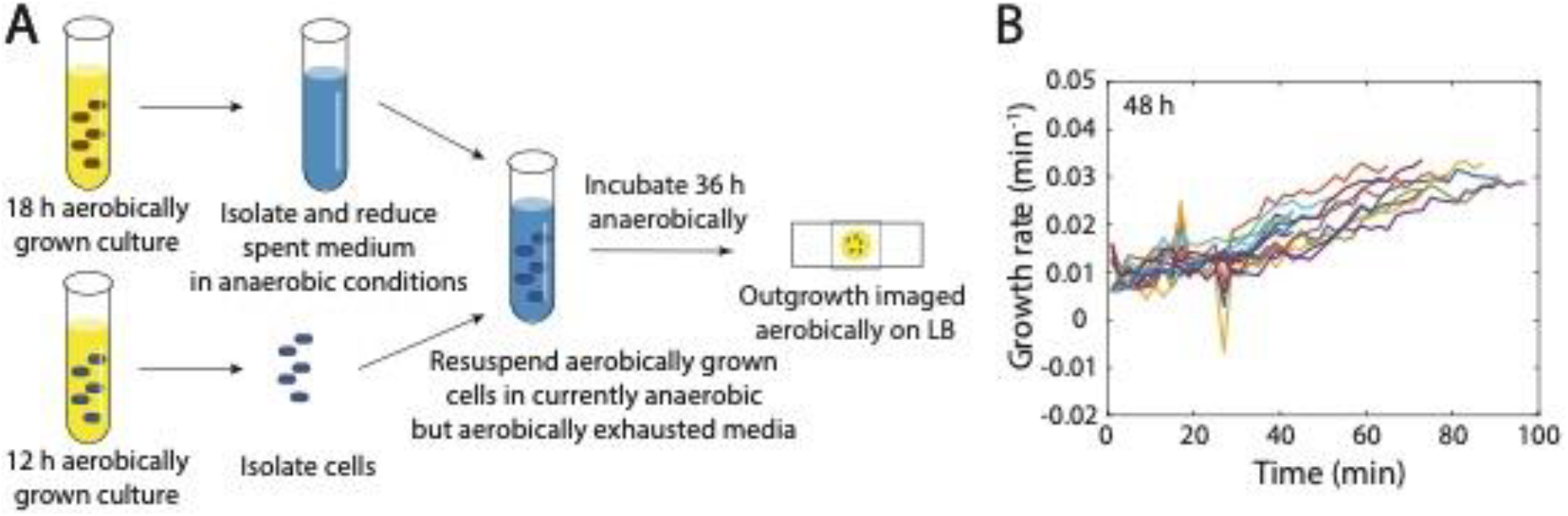
Anaerobic incubation in stationary phase prevents dormancy. A) A 12-h aerobically grown culture entering stationary phase was resuspended in the supernatant from an 18-h aerobically grown culture pre-reduced in an anaerobic chamber. The resulting culture was incubated in an anaerobic chamber for 36 h, and then outgrowth was monitored aerobically on fresh LB agarose pads. B) After the protocol in (A), all cells grew immediately and accelerated in growth similar to aerobically grown cultures after 12 h of incubation (Fig. 1B).

## Discussion

Here, we discovered that dilution of an aerobically grown stationary-phase *E. coli* culture into fresh medium led to a heterogeneous population of immediately growing, delayed-growth, and non-growing cells (Fig. 1). Longer times spent in stationary phase led to an increased fraction of non-growing cells, although a small fraction remained able to grow immediately (Fig. 1K). Dormant cells exhibited bright foci that dissolved prior to the resumption of growth (Fig. 3B) and after the synthesis of repair enzymes (Fig. 5C-F, S2A,B), supporting the hypothesis that dormancy is due to intracellular protein and DNA damage. When cells were grown in conditions that precluded respiration, dormancy did not occur and bright foci could not be detected (Fig. 7B). Mathematical modeling based on gradual damage accumulation and division-mediated recovery through asymmetric partitioning of damage quantitatively predicted the fraction of each of the three subpopulations over time (Fig. 6D), with maintenance of an immediately growing fraction of ~10%. Together, the evidence indicates that dormancy is driven by the accumulation of damage related to respiration. This model should prove powerful for interpreting the molecular basis of dormancy in other strains and species.

The ubiquity of dormancy across natural isolates (Fig. 1M,N) raises the question of whether delayed growth is an inevitable effect of damage or a programmed response. Several lines of evidence suggest the latter. Long lag times are beneficial for survival of antibiotic treatment (24) and likely other stresses, despite their obvious disadvantages in conditions that permit rapid growth. Moreover, it is possible to select for mutants with increased lag and increased persistence (35), underscoring the genetic potential for increasing lag under certain selective pressures. Evolution experiments with repeated passaging have selected for shorter lag times due to competition for nutrients (30), and these strains may as a result have reduced fitness in LB in long-term stationary phase. Although cell size is highly correlated with lag time in some conditions (30), the mechanism connecting the two has not been elucidated. Nevertheless, our discovery that bigger cells are less likely to become dormant (Fig. 4) suggests a direct connection between size and stationary/lag phase. Dormancy may thus act as a selective pressure against cells becoming too small (36). Regardless, our studies underscore the dependence of stationary-phase physiology on how cells enter starvation, and provide a road map for studying dormancy across mutants and nutrient environments.

Our data extends a growing, common framework for growth-deficient cells, linking stationary phase to persistence and aging through protein aggregation, delayed growth, and oxygen dependence. The nature and dynamics of aggregate formation remain a substantial mystery. The fact that aggregates are comparable in size to the diffraction limit in delayed-growth cells suggests that they form earlier in stationary phase but are too small to resolve. Until they reach a sufficiently large size (similar to that of polysomes), they may not end up segregated to the poles (37), and if they are not at the poles, they may not be asymmetrically segregated upon division. Aging studies generally focus on exponential growth, although there are physiological similarities between the aggregates that accumulate over generations of steady-state growth and in stationary phase. Our data suggest that aging would be accelerated if cells switched between exponential and stationary phase due to starvation-induced damage accumulation.

The low but nonzero level of persister formation during exponential growth (38) and the small growth defect of cells with older poles (29) suggests that damage is a small but ever-present challenge for bacteria. The same damage that leads to dormancy may also be the cause of the variability in single-cell exponential growth rates (39), and in any event will introduce variability into any studies of stationary-phase outgrowth (12). The dissolution of aggregates releases many factors important for growth, and their sequestration could act as a transcriptional signal in stationary phase (29). Although the molecular triggers for aggregate formation have yet to be determined, our data suggest that respiration plays a major role (Fig. 6B), which implies that facultative anaerobes may generally exhibit more growth heterogeneity in aerobic conditions. The ultimate consequences of cellular inability to remove aggregates are manifested in stationary phase, evidenced by the aberrant morphologies and regrowth behaviors of Δ*dnaK* cells (Fig. 5H-J) despite growth that is similar to wild-type during log and early stationary phase (Fig. 5G).

It is likely that most bacteria frequently enter stationary phase in their natural environment, particularly in the gut where periodic feeding leads to feast and famine. A screen of *B. subtilis* CRISPRi knockdowns suggested that levels of essential genes have been determined by the need to survive stationary phase rather than for rapid growth (41), indicating that stationary phase has proven a strong selective pressure. Indeed, a recent laboratory evolution experiment identified a Pareto front (situation in which improving one phenotype necessitates negative effects on other phenotypes) of adaptive mutations with tradeoffs between respiration and stationary phase in budding yeast (42). Moreover, daily passaging of *E. coli* selected for mutations that reduce fitness in long-term stationary phase (43). The heterogeneity of dormancy naturally presents the possibility for bet hedging, and similar heterogeneity could arise from other stresses that cause damage. Future investigation of species from environments such as the soil that constantly experience fluctuations in oxidative stress, compared with gut commensals (most of which are obligate anaerobes), may help to shed light on selection in stationary phase. Ultimately, stationary phase presents fascinating physiological challenges that provide a window into the fundamental mechanisms of growth.

## Author Contributions

S.C., L.W., and K.C.H. designed the research; S.C. and L.W. performed the research; S.C., L.W., and K.C.H. analyzed the data; and S.C., L.W., and K.C.H wrote the paper.

## Acknowledgements

The authors thank Manohary Rajendram, Handuo Shi, and members of the Huang lab for helpful discussions. The authors acknowledge funding from a Stanford Graduate Fellowship and a National Science Foundation Graduate Research Fellowship (to. S.C.) and from the Allen Discovery Center at Stanford on Systems Modeling of Infection (to K.C.H). K.C.H. is a Chan Zuckerberg Biohub Investigator.

## Methods

### Strain culturing

Strains used in this study can be found in Table S1. All cultures were grown in 3 mL of filter-sterilized LB at 37 °C in glass test tubes with constant shaking in aerobic conditions. Cultures were started at an OD at 600 nm (OD_600_) of 0.1 with cells from a 5-7 h culture inoculated from a frozen stock. Spent medium was isolated by spinning down (at 4000*g* for 5 min) and filtering a saturated culture with a 0.22-μm polyethersulfone filter (Millex-GP SLGP033RS).

### Measurement of population growth metrics

Growth curves were obtained using an Epoch 2 Microplate Spectrophotometer (Biotek Instruments, Vermont). The plate reader went through 15-min cycles of incubation at 37 °C, shaking linearly for 145 s, and then absorbance measurements (wavelength 600 nm, 25 flashes, 2-ms settle between flashes).

### Single-cell imaging

One microliter of cells was diluted 1:200 with fresh medium, spotted onto a pad of 1% agarose+LB, and imaged on a Nikon Eclipse Ti-E inverted fluorescence microscope with a 100X (NA 1.40) oil-immersion objective (Nikon Instruments). Phase-contrast and epifluorescence images were collected on a DU885 electron-multiplying CCD camera (Andor Technology) or a Neo sCMOS camera (Andor Technology) using μManager v. 1.4 (44). Cells were maintained at 37 °C during imaging with an active-control environmental chamber (Haison Technology).

### Image analysis

The MATLAB (MathWorks, Natick, MA, USA) image processing code *Morphometrics* was used to segment cells and to identify cell outlines from phase-contrast microscopy images. A local coordinate system was generated for each cell outline using a method adapted from *MicrobeTracker* (46). Cell widths were calculated by averaging the distances between contour points perpendicular to the cell midline, excluding contour points within the poles and sites of septation. Cell length was calculated as the length of the midline from pole to pole. Cell area was used to calculate growth rate. Cell volume was estimated from cylindrical surfaces of revolution of local width measurements. Cellular dimensions (width/length/area/volume) were quantified by averaging single-cell results across a population. See figure legends for the number of cells analyzed (*n*) and error bar definitions.

### Classification of subpopulations

A cell was classified as immediately growing if its median growth rate in the first 20 min of imaging was >0.004 min^−1^ and its growth rate eventually exceeded 0.01 min^−1^. Otherwise, a cell was classified as delayed growth if its growth rate exceeded >0.005 min^−1^. The remaining cells were classified as nongrowing if their average growth rate at the conclusion of imaging was <0.003 min^−1^.

### Detection of intracellular aggregates

Bright spots within cells in phase-contrast images were identified as aggregates if their maximum intensity was >5000 a.u. brighter than the median brightness of all pixels within the cell that were >2 pixels from the perimeter.

### Deep learning classification of single-cell images

Densenet121 (47) was trained on labeled images of cells cropped to 64×64 pixels. The training set included 2333 images of immediately growing cells, 788 images of delayed-growth cells, and 3225 images of non-growing cells, from 9 imaging experiments. To augment the dataset, all images were mirrored and rotated 90, 180, and 270 degrees, resulting in 7 additional images per original image. Images were randomly partitioned between the training, development, and test datasets with an 80%/10%/10% breakdown. All augmented images were placed in the same dataset as the original image. L2 regularization was added to Densenet121 with a value of 0.0001. The parameters used for analysis of the test set were those from the 22^nd^ epoch of training, which was selected based on its performance with the development set.

### Mathematical model and simulations

In our model (Fig. 6, S3), throughout stationary phase starting from ~11 h after inoculation, cells accumulate damage from an initial value of 0, divide with a rate *ρ*, and transition between regrowth states (immediately growing, delayed growth, non-growing over the period of observation, and non-viable) according to their damage levels. In a short time Δ*t*, a cell with damage *A* undergoes the following: 1) damage increases to *A* + 1 with probability δΔ*t*; 2) division occurs with probability *ρ*Δ*t*; and 3) if *A* > *A*_2_, the cell becomes nonviable with probability *γ*Δ*t*. During growth on fresh medium, cells are classified as immediately growing if *A* ≤ *A*_1_, delayed-growth if *A*_1_ < *A* ≤ *A*_2_, and non-growing if *A* > *A*_2_.

Upon division, damage partitioning is asymmetric: one daughter receives all damage, while the other daughter is returned to an undamaged state. Division always coincides with the removal of a randomly selected cell to maintain population size. Simulations based on this model were carried out using the Gillespie Algorithm (48) with 1,000 cells. Analytical expressions were derived from an ODE continuum model that matches simulations of the probabilistic model in the long-time limit (Supplementary Text). A generalization in which the division rate *ρ* decreases as damage increases is discussed in the Supplementary Text.

### Data availability

All data used in this manuscript are growth curves and microscopy images. All data and associated software are available upon request from the corresponding author.

## Supplementary Text

### Analytical expressions for the proportion of immediately growing and dormant cells

To derive analytical expressions for the steady states of our model, we specify a continuum version of our stochastic model of damage-induced dormancy that has the same steady states as the stochastic model. In the continuum model, *g* is the fraction of immediately growing cells with aggregates *A* < *A*_1_, *r* is the fraction of delayed cells with *A*_1_ < *A* < *A*_2_, 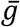 is the fraction of cells with *A* > *A*_2_ that do not grow but still divide, and *e* is the fraction of cells that cannot divide or that are dead but not lysed. Average deterministic equations corresponding to the stochastic model are:

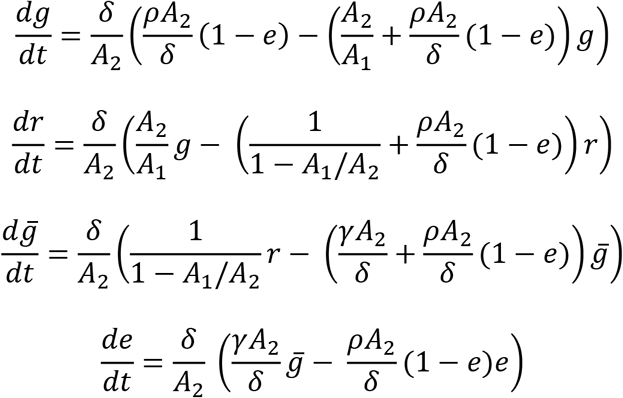

where we select 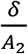 to govern the timescale, and other governing parameters are *A*_1_/*A*_2_, 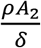, and 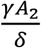. For model (b) with damage-induced dormancy (δ > 0) and division (*ρ* > 0) but no death (*γ* = 0, *e* = 0), at steady state 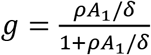 and 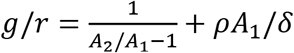. For model (c) including death (*γ* > 0), all cells eventually die (so eventually *e* = 1) when *ρ*^−1^ > *A*_2_δ^−1^ + *γ*^−1^, which is the case for the parameters in Fig. 6D. Fig. S3B shows how steady states but not earlier dynamics of the stochastic model match the continuum model.

### Dynamics are similar when the division rate decreases linearly with damage

Here, instead of a constant division rate *ρ* and a constant rate *γ* of switching to non-division (or death without lysis) when *A* ≥ *A*_2_ in model (c) (Fig. 6A), division rate decreases with damage according to 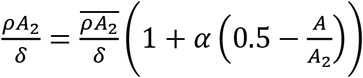 and 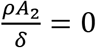 if *A* ≥ *A*_2_(α^−1^ + 0.5). Thus, when *A* = *A*_2_(α^−1^ + 0.5), cells enter a non-dividing state. If α = 0, the non-dividing state is never reached, whereas if α = 2, division halts when *A* = *A*_2_ coincidentally with the cell entering the non-growing state.

## Supplemental Figure Legends

**Figure S1:**
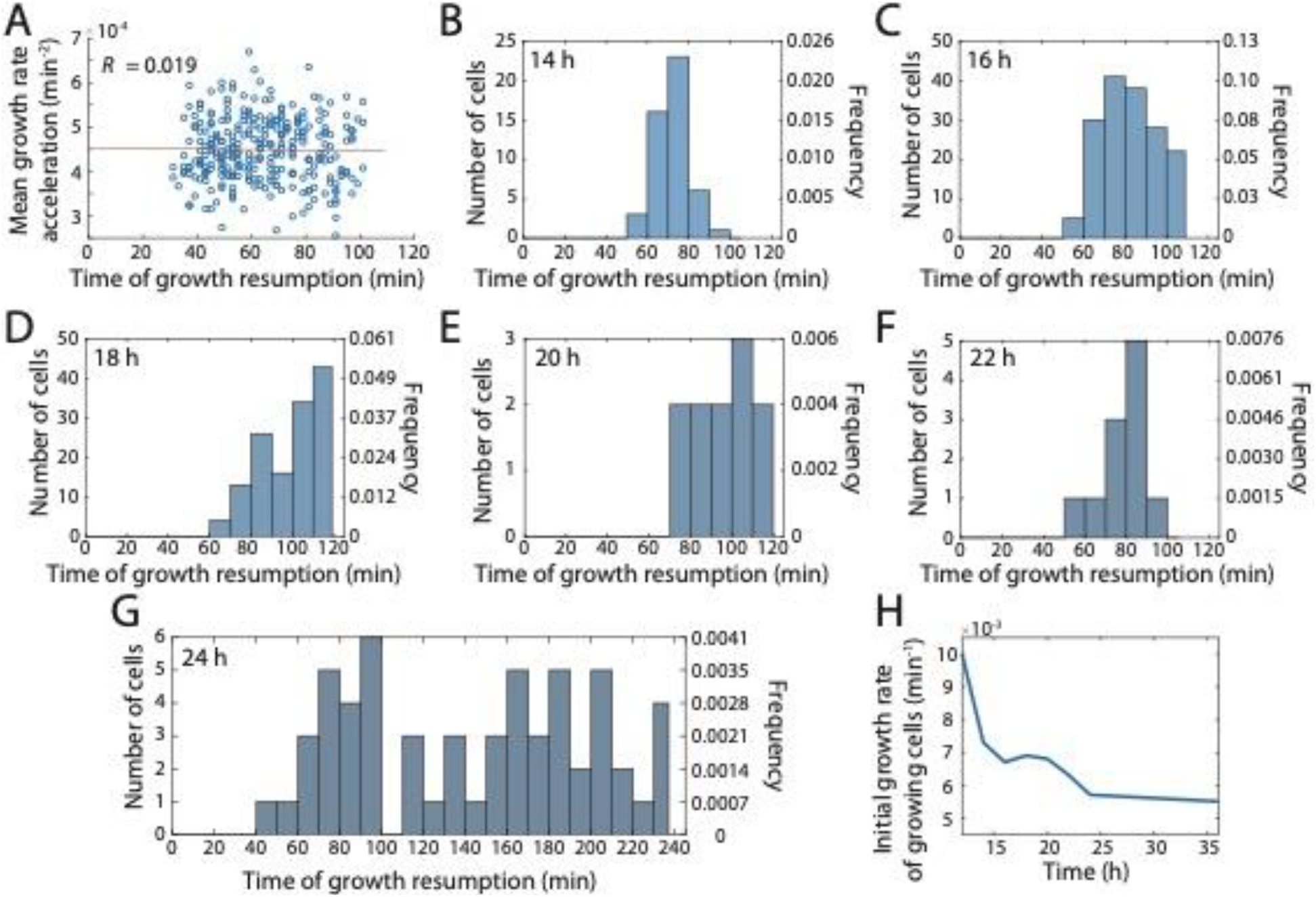
The time at which cells resume growth increases with increasing time in stationary phase, and immediately growing cells start with a lower growth rate. A) The average acceleration in growth rate (computed for each cell as the slope of a line fitted to data from 18 min before to 8 min after achieving a growth rate of 0.02 min^−1^) was uncorrelated with the time at which cells from a 16-h culture resumed growth. B-G) The time at which cells resumed growth increased with increasing culture age. Dashed lines are means. The frequency was measured including non-growing cells. H) The initial growth rate of immediately growing cells decreased as a function of culture age.

**Figure S2:**
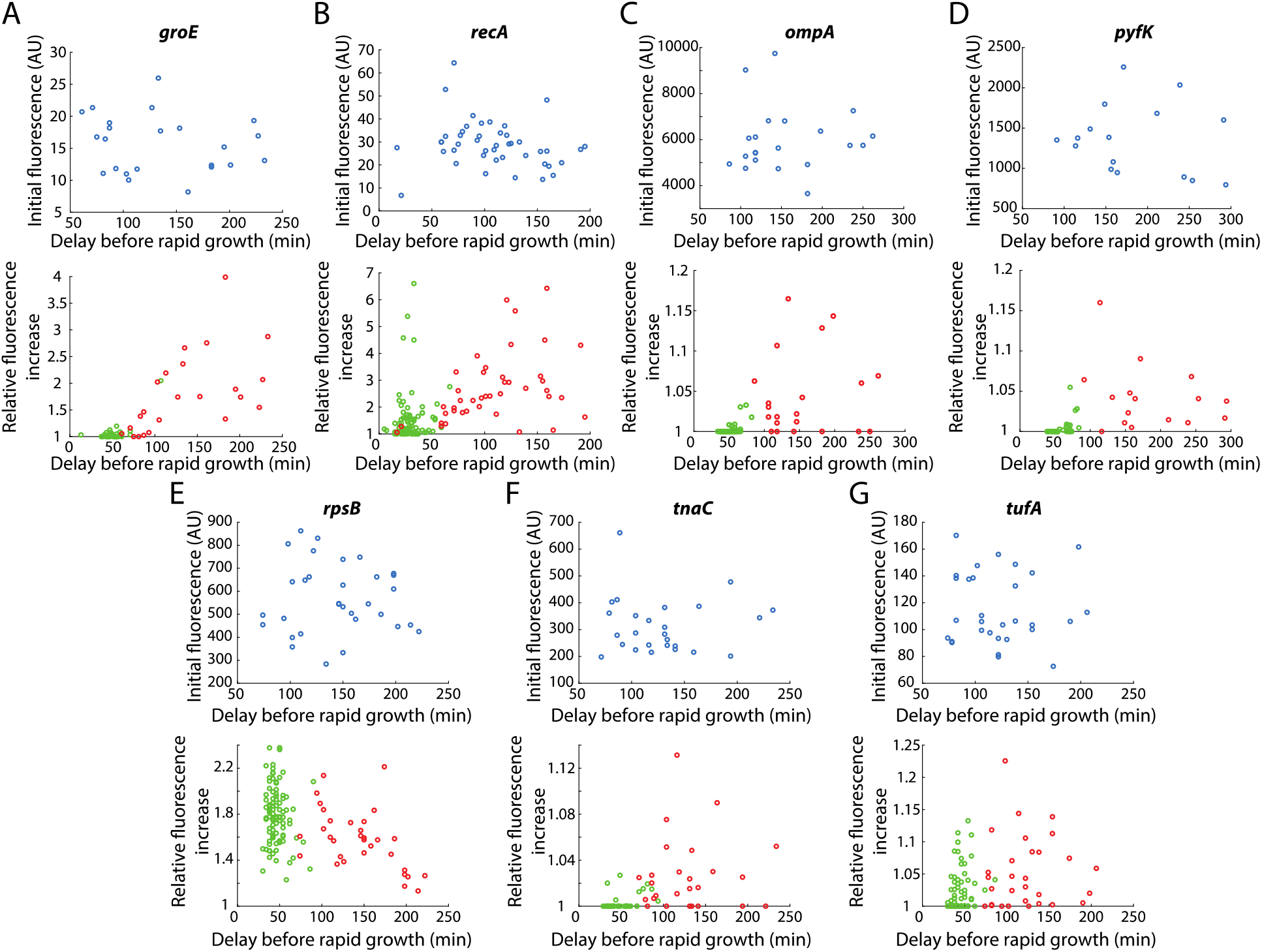
Repair enzymes are upregulated in delayed growth cells, but not genes related to metabolism or translation. A,B) Similar to *dnaK* (Fig. 5B,E), initial GFP fluorescence from the promoter of *groE* (A) or *recA* (B) was uncorrelated with the delay before rapid growth (defined as the time to reach growth rate 0.02 min^−1^) (top), but the maximum fluorescence during time-lapse imaging on fresh medium relative to the initial fluorescence was correlated with the delay before rapid growth (bottom). These data support the hypothesis that dormancy is due to accumulated DNA and protein damage. C-G) By contrast, for several non-damage-related promoters including *ompA* (C), *pyfK* (D), *rpsB* (E), *tnaC* (F), and *tufA* (G), GFP increases were not correlated with the delay until rapid growth.

**Figure S3:**
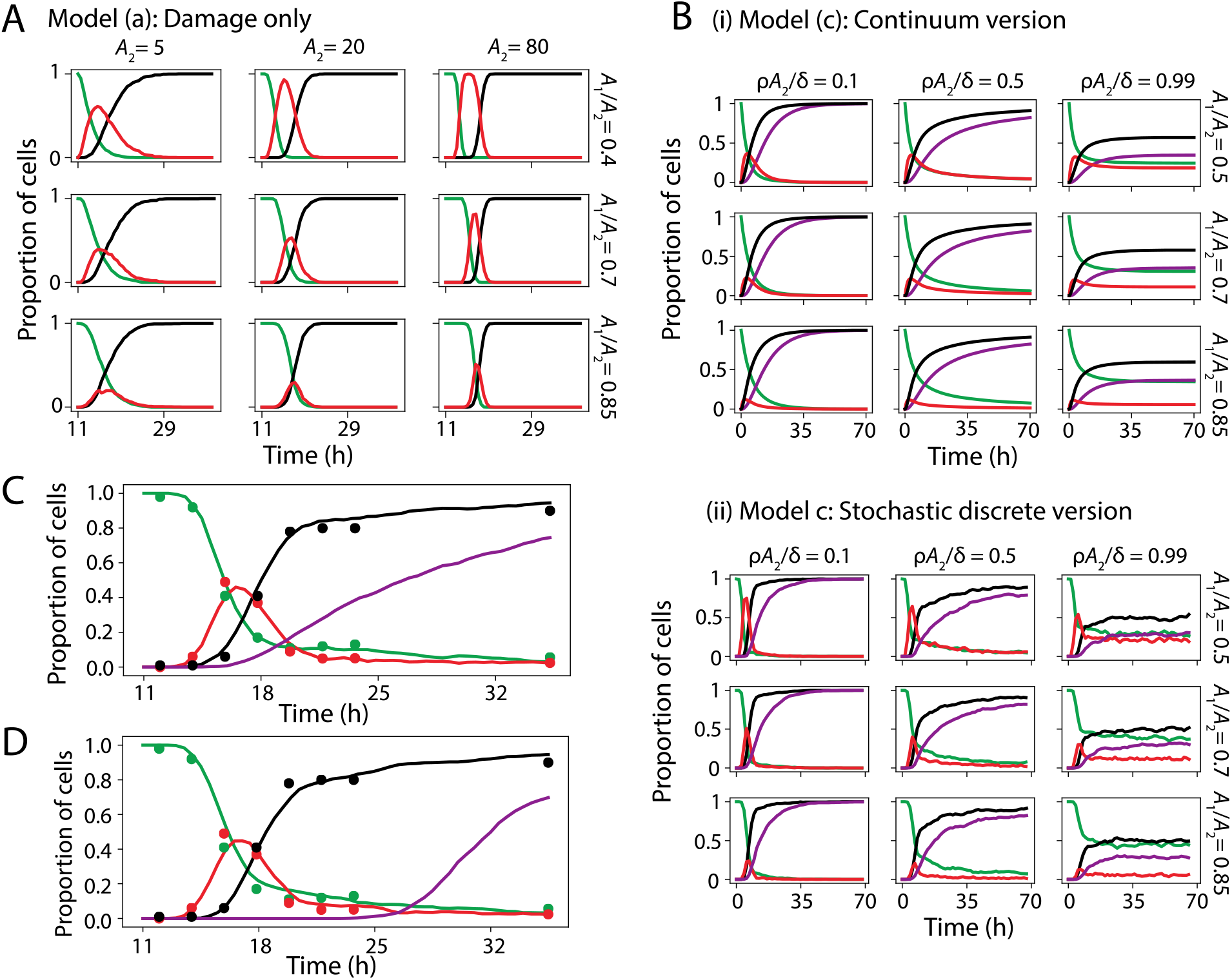
Mathematical model makes similar predictions if a threshold intracellular concentration of damage triggers dormancy, or if division rate declines with damage. (A) Simulations of model (a) (Fig. 6A) with damage-induced dormancy but no division or death recapitulated regrowth statistics up to, but not beyond, ~20 h after inoculation when 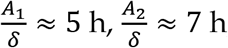, so 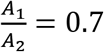, and *A*_2_ = 20. In all panels, immediately growing cells are green, dormant cells are red, and non-growing cells are black. *A*_2_ tuned the time of the peak in the delayed-regrowth fraction, while both the peak and duration of the delayed regrowth fraction were tuned by *A*_1_/*A*_2_. (B) Steady states but not the early dynamics of the continuum model (i) (Supplementary Text) agreed with the stochastic model (ii), unless *A*_2_ ≈ 1 or *A*_1_ ≈ *A*_2_ in which case discretization effects were strong (compare bottom right panels for (i) and (ii)). Parameters are 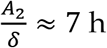, *A* = 20 (for (i)), and 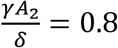, while 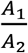 and 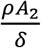 vary. Purple indicates the subpopulation of non-dividing or dead cells. (C) The dynamics of model (c) are similar if a threshold intracellular concentration of damage triggers dormancy, as opposed to a threshold absolute level of damage. Here, *A* now represents damage concentration, which doubles in the daughter cell that inherits all damage upon division. Other parameter definitions are unchanged. Parameters 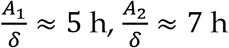, so 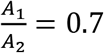, *A*_2_ = 20, 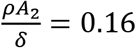, and 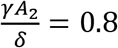, as for Fig. 6D, produce an excellent fit. Solid lines correspond to the model and circles correspond to data. (D) The dynamics of model (c) are similar if instead of a death rate *γ* > 0 when *A* > *A*_2_, the division rate declines linearly to 0 with increased damage (Supplementary Text). Parameters 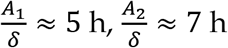, so 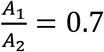, *A*_2_ = 20, 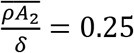, and α = 0.45 produce an excellent fit with a slight lag in cell death (purple). Solid lines correspond to the model and circles correspond to data.

**Figure S4:**
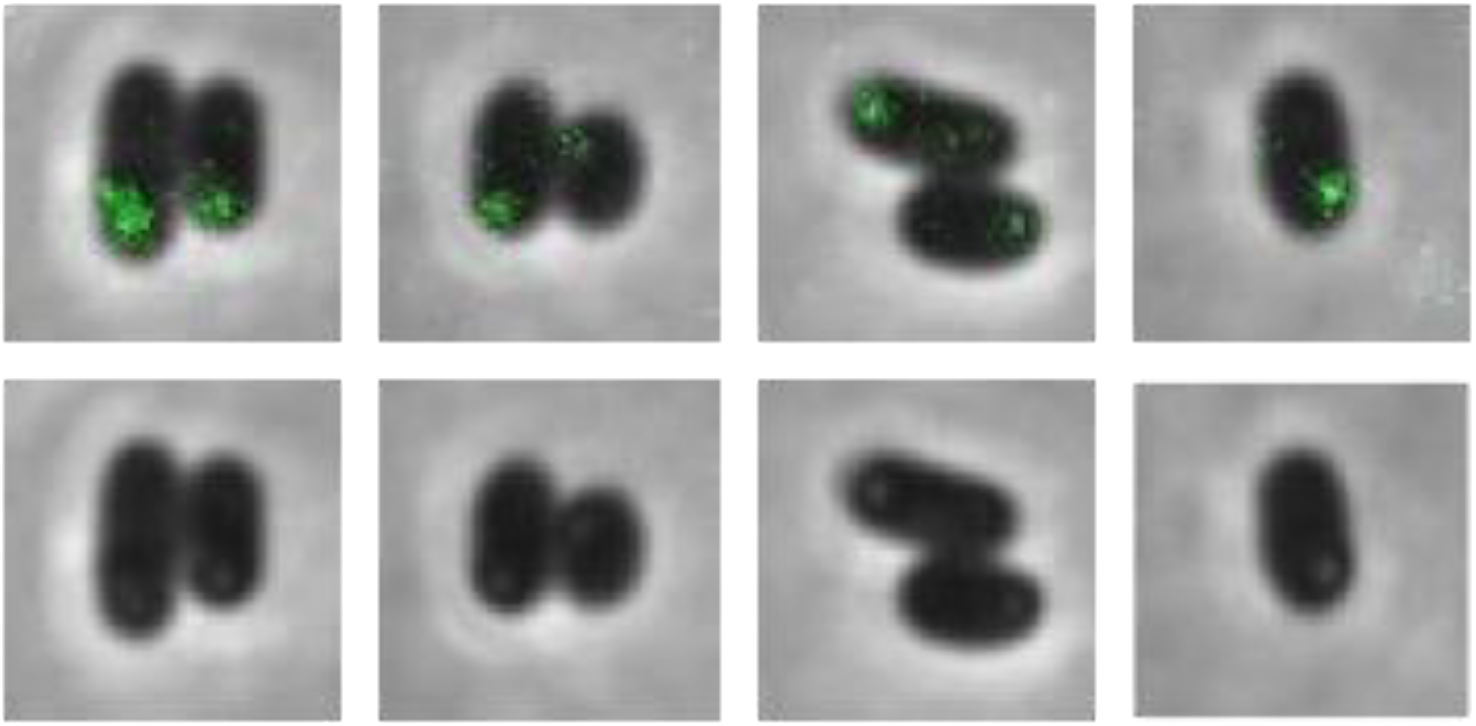
FtsZ is sequestered in aggregates at the cell poles in stationary phase. Top: overlay of FtsZ-msfGFP fluorescence with phase-contrast images. Bottom: phase-contrast images showing bright foci at the same intracellular locations as the FtsZ punctae.

**Figure S5:**
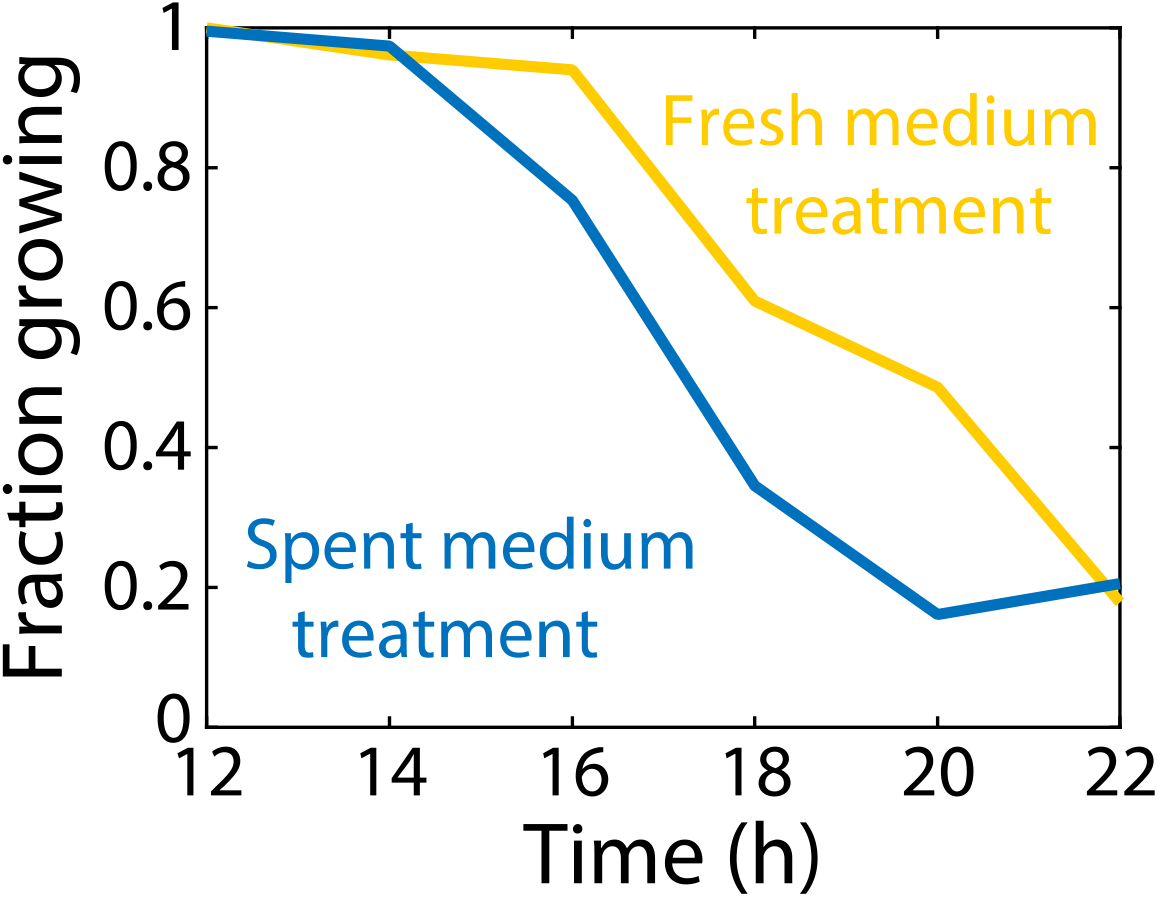
A pulse of fresh medium delays the onset of dormancy. Resuspension of cells from a 12-h culture in fresh LB for 20 min (yellow) led to a ~2-h delay in the onset of dormancy following resuspension in spent supernatant relative to a control that did not experience the pulse of fresh medium, indicating that the cause of dormancy can be reversed.

**Figure S6:**
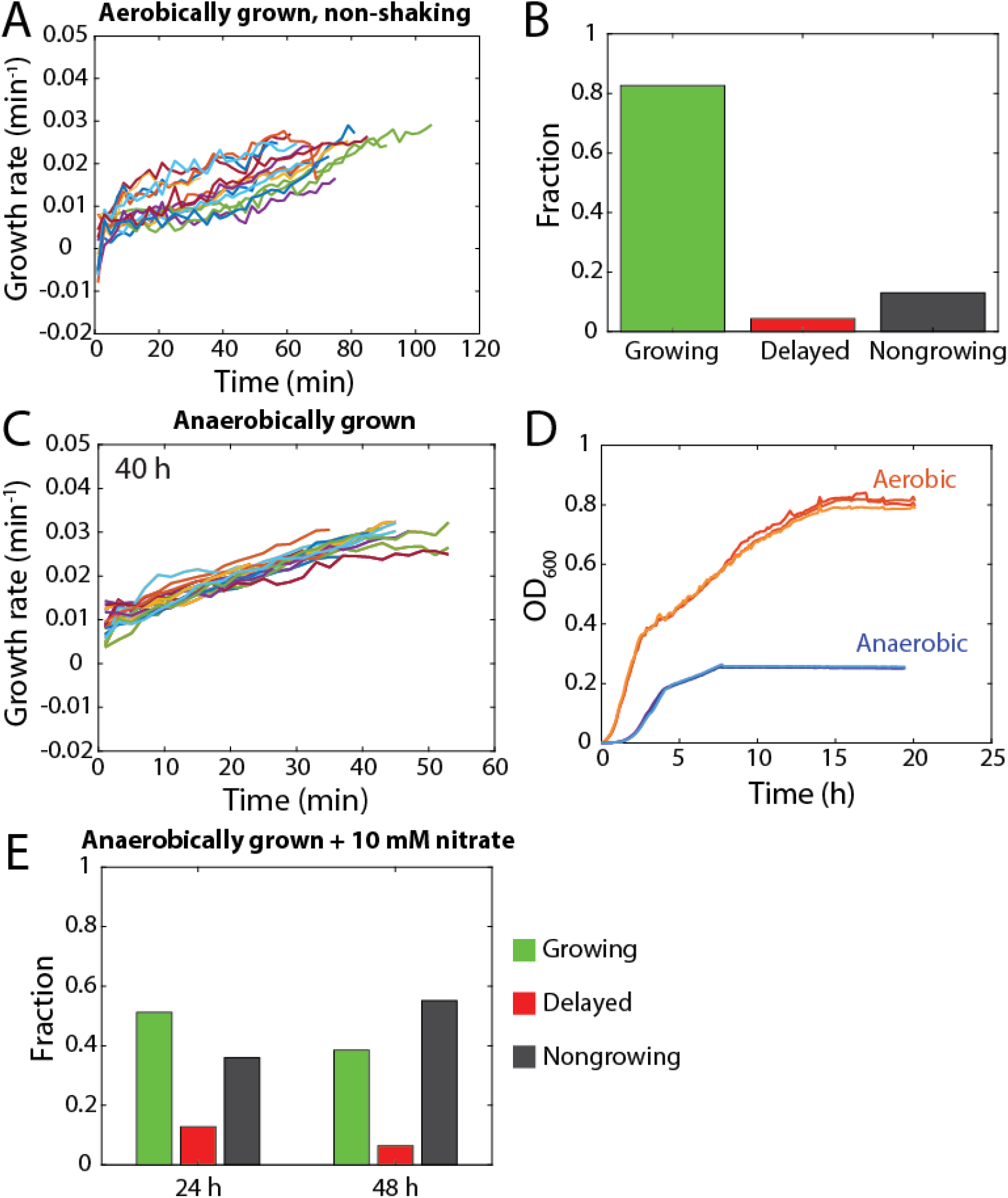
Anaerobically grown cells resume growth immediately coming from stationary phase unless nitrate is available for respiration during stationary phase. A) After 20 h of growth in non-shaking conditions, most cells were able to grow immediately upon exposure to fresh medium, unlike in shaking conditions (Fig. 1K). B) Fractions of each subpopulation after 20 h of aerobic growth in non-shaking conditions show that most cells can grow immediately. C) Cells grown in an anerobic chamber for 40 h immediately resumed growth on fresh LB agarose pads, suggesting oxidative stress is a primary cause of dormancy. D) *E. coli* saturates at a lower yield in LB when grown in anaerobic versus aerobic conditions. E) After 24 h of anaerobic growth in LB supplemented with 10 mM nitrate, almost half of the population was dormant or nongrowing. After 48 h, the nongrowing population expanded at the expense of the fractions of both immediately growing and dormant cells.

## Supplemental Movie Legends

**Movie S1: Some Δ*dnaK* cells exhibit cycles of growth and shrinking during emergence from stationary phase.**

Upper left: the time since being exposed to fresh medium, in hours and minutes.

